# Robotic Multi-Probe-Single-Actuator Inchworm Neural Microdrive

**DOI:** 10.1101/2020.10.13.338137

**Authors:** R. D. Smith, I. Kolb, S. Tanaka, A. K. Lee, T. D. Harris, M. Barbic

## Abstract

Electrophysiology is one of the major experimental techniques used in neuroscience. The favorable spatial and temporal resolution as well as the increasingly larger site counts of brain recording electrodes contribute to the popularity and importance of electrophysiology in neuroscience. Such electrodes are typically mechanically placed in the brain to perform acute or chronic freely moving animal measurements. The micro positioners currently used for such tasks employ a single translator per independent probe being placed into the targeted brain region, leading to significant size and weight restrictions. To overcome this limitation, we have developed a miniature robotic multi-probe neural microdrive that utilizes novel phase-change-material-filled resistive heater micro-grippers. The microscopic dimensions, gentle gripping action, independent electronic actuation control, and high packing density of the grippers allow for micrometer-precision independent positioning of multiple arbitrarily shaped parallel neural electrodes with only a single piezo actuator in an inchworm motor configuration. This multi-probe-single-actuator design allows for significant size and weight reduction, as well as remote control and potential automation of the microdrive. We demonstrate accurate placement of multiple independent recording electrodes into the CA1 region of the rat hippocampus *in vivo* in acute and chronic settings. Thus, our robotic neural microdrive technology is applicable towards basic neuroscience and clinical studies, as well as other multi-probe or multi-sensor micro-positioning applications.

**One Sentence Summary:** Miniature robotic multi-probe single-actuator microdrive utilizing phase change material based micro-grippers.

## Introduction

The miniaturization of mechanical, electrical, optical and other tools and devices has been steadily ongoing for decades, enabled by advances in manufacturing, electronics, and novel mechanical designs. For scientific instruments, miniaturization is often motivated by requirements for enhancing sensitivity and functionality, reducing costs, parallelizing and scaling of experiments, and demands for weight and size reduction to enable operation in constrained spaces. Aided by similar technological advances, tools and techniques in neuroscience have undergone tremendous progress, enabling the study of the living, behaving brain^1–7^.

A variety of electrical, chemical, and optical probes are mechanically inserted into the brain to perform in vivo measurements or perturbations on acutely or chronically implanted animals^8–24^. Since the brain is anatomically structured into different functional regions, these probes must be precisely placed in the region of scientific or clinical interest, often with micron-scale accuracy. A number of microfabricated electrodes have enabled recording from neurons localized to a single plane^25^ or column^22^, but for many applications multiple independently actuated electrodes are required to record neural activity in a geometry-flexible fashion. Therefore, a mechanical positioner, commonly termed microdrive, is usually employed for such adjustable placement tasks and its design has undergone steady technological development and improvement over the years^26–46^. Presently available neural probe microdrives tend to be manually operated devices where each independent probe is assigned its own lead-screw-based positioner. Increasing the number of neural probes mounted on such drives often makes them complex, bulky, and heavy, and the practice of the probe placement becomes increasingly cumbersome, time consuming, and disruptive to the scientific experiment. Some effort has gone into developing motorized microdrive devices that would allow for more efficient electrophysiology data collection through parallelization and remote computer-controlled operation without human intervention^47–60^. However, even in these designs, each independent neural probe is assigned its own independent motorized positioner which often makes the size, weight, complexity, and expense of the microdrive prohibitive.

Here we demonstrate a new design of a robotic inchworm neural microdrive that utilizes novel electronically controlled and densely packed probe micro-grippers in combination with only a single piezo actuator for independent translation of each loaded neural probe. The multi-probe-single-actuator (MPSA) concept and design therefore allows for a significant reduction in complexity, size, weight, and cost, while still providing micron-scale independent positioning control of each neural electrode. As a proof of concept, we constructed and demonstrated the operation of the microdrive while loaded with classic twisted wire tetrode neural electrodes. We performed remote-controlled placement of these independent recording neural electrodes with micrometer precision into the CA1 region of the rat hippocampus *in vivo* in acute and chronic settings.

## Results

### Phase-change-material-filled resistive heater coil micro-gripper

We introduce the new probe micro-gripper in Figure 1 A-B, which schematically describes the functional design of the device. A helical resistive wire coil embedded in a printed circuit board (PCB) non-plated via is connected such that an electric current from a power source can be passed through the coil. This allows for the electronically controlled change of the gripper’s temperature through Joule heating. A probe is threaded through the bore of the coil and the remainder is filled with a temperature controlled phase change material (PCM)^61^. When the heater coil does not carry current (Figure 1A), the gripper is at the ambient temperature, the PCM in the bore is in the solid state, and therefore the neural probe is “gripped”^61^. When the heater coil carries current (Figure 1B), the gripper heats up, the PCM in the bore melts and goes into the liquid state. The neural probe is therefore “released”^61^ and can be moved axially through the gripper. Critically, due to the microscopic dimensions of the space between the gripper and the neural probe, the PCM can be maintained in the bore through capillary action without exiting during probe motion. Figure 1C shows a photograph of the side view of the fabricated helical resistive micro-heater before it is placed into a PCB, while Figure 1D shows a photograph of the top view of the heater coil after it is installed (see Materials and Methods).

**Figure 1.**
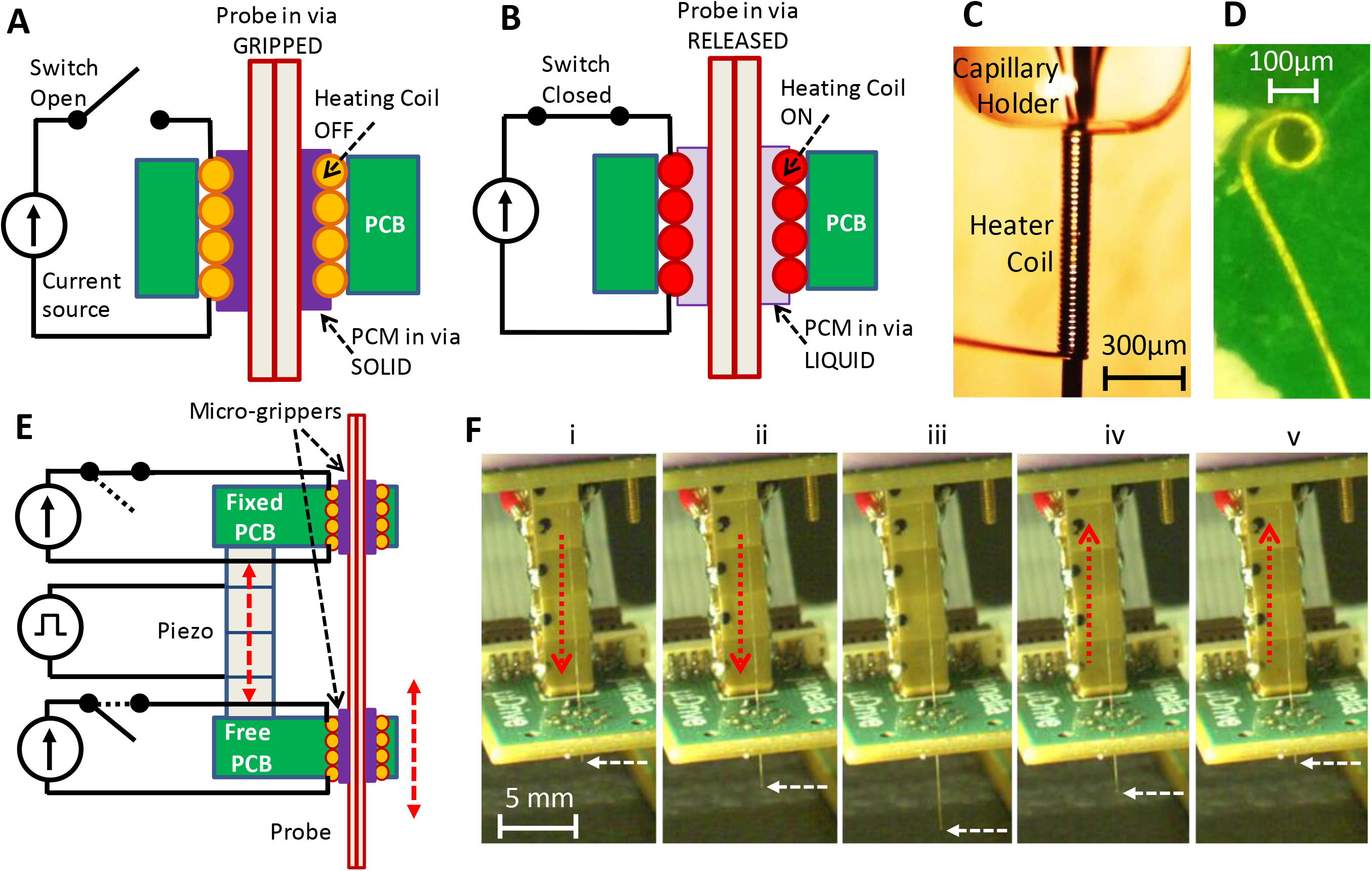
Micro-gripper and single probe inchworm microdrive design. (A-B) Side view of micro-gripper design. A resistive heating coil is installed in a printed circuit board via. A probe is placed through the coil bore and the bore is filled with a temperature-responsive phase change material (PCM). With the heating coil off (A), PCM is in the solid state and the probe is “gripped”. When the coil is on (B), PCM is in the liquid state and the probe in the micro-gripper is “released” and can be moved axially through the bore. The PCM can be maintained in the micro-gripper through capillary action and does not exit the bore. (C) Photograph of the fabricated heating coil before it is placed into the board. (D) Photograph of the top view of the heating coil installed into the printed circuit board. (E) Schematic diagram of the inchworm motor design. Two parallel PCBs, eachwith the installed micro-grippers are aligned along the probe axis and joined by a single piezo actuator. The top board is fixed while the bottom board is movable by the motion of the voltage-controlled piezo actuator. Sequential electronically-controlled probe gripping and piezo extension and contraction (described in Figure 2) are used for probe translation (white dashed arrow points to the probe tip) in either direction (up or down), as shown in F (i)-(v) and Supplementary Video 1.

We emphasize several important features of this probe gripper design. First, the gripper is operated electronically which allows for remote control. Second, the gripper is dimensionally microscopic (~75μm inner diameter/110μm outer diameter, Figure 1 C-D) and of similar cross-sectional size as the neural probe that it grips (twisted wire tetrode probe ~55μm diameter). Therefore, the gripper takes up minimal PCB area, and many probes can be densely packed, as we describe in a later section. Third, the probe gripped within the bore of the device can be of any cross-sectional shape, as the liquid PCM in the released state conforms to the probe shape before it solidifies into the gripped state. This presents an opportunity to use the device to grip a variety of probes such as electrical testing probes, optical fibers, silicon neural electrodes, glass pipettes/capillary probes, carbon fibers, and ultrasonic probes. For the purposes of demonstration, in addition to neural twisted wire tetrode electrodes, we also demonstrate in a later section gripping and translation of glass micropipettes.

### Single probe inchworm microdrive operation

The above described neural probe micro-gripper design is integrated into an inchworm motor structure, as shown diagrammatically in Figure 1E. The motor is constructed from two parallel PCBs, each with the embedded helical coil heater grippers aligned along the neural probe axis. The two parallel boards are joined by a single piezoelectric block stack actuator that is controlled by a piezo actuator voltage driver (see Materials and Methods). In this diagram, the top board is fixed while the bottom board is free and movable by the motion of the piezo actuator. The functional concept will remain the same if the roles are reversed so that the choice can be dictated by application considerations. The inchworm motor steps^62,63^ of sequential probe gripping and piezo extension and contraction (as we describe in detail in the next section) are used for the neural probe translation shown in Figure 1F. By electronically controlling the current though the resistive heaters (and therefore the gripping and releasing of the probe in the bores) in the respective top and bottom boards (Figure 1E), the neural probe in the device can be translated in either direction (up or down) with the piezo actuator, as shown in Figure 1F and Supplementary Video 1.

We note several important features of this inchworm motor design. First, the piezo actuator is, just like the probe grippers, controlled electronically which allows for remote control. Second, the piezo actuator has micron-scale translation capability, accuracy, and repeatability. This is critical when precise positioning of the neural electrodes is required for targeted neural recordings. Third, the inchworm motor design in general allows for the long range forward or backward motion of the probe with indefinite translation range. This is because the piezo can make a practically unlimited number of microscopic steps, as Figure 1F illustrates.

We now provide detailed description, in Figure 2, of the sequence of electronic actuation signals sent to the inchworm microdrive for its three distinct probe translation operations: (a) downward translation step, (b) upward translation step, and (c) no translation step. Translation of a probe is accomplished using a precisely timed sequence of electronic actuation steps (Figure 2). The sequence to move a probe one step downwards is depicted in Figure 2A. In the first step of the sequence, Figure 2A(i), the top heating coil is turned on by passing a controlled current through it. This causes the temperature of the top micro-gripper to rise, such that the PCM in the top heater bore melts and releases the probe. While the top micro-gripper’s PCM is liquid, a voltage is applied to the piezo actuator, causing it to extend downward, as shown in Figure 2A(ii). Since the probe is still gripped in the bottom board, which moves with the piezo actuator, the probe is also moved downward the same distance, sliding through the top micro-gripper. After the top heater is turned off, the top PCM remains liquid for a short time while the heat diffuses away. Subsequently, the PCM solidifies and the top micro-gripper re-grips the probe while, importantly, the piezo actuator is still extended, as shown in Figure 2A(iii). Next, the bottom heating coil is turned on, which causes the bottom micro-gripper to release the probe, as shown in Figure 2A(iv). While the bottom PCM is liquid, the voltage applied to the piezo actuator is set to zero, causing the piezo to contract upward, back to its initial length, as shown in Figure 2A(v). Since the probe is released in the bottom board (which moves with the piezo actuator) and is still gripped in the stationary top board, the probe remains fixed and does not move back with the piezo actuator, as shown in Figure 2A(v). After a period with the bottom heater off, the bottom PCM solidifies, and the bottom micro-gripper re-grips the probe, as shown in Figure 2A(vi). The result of this sequence of six specifically timed steps is a single downward translation step of the probe with a step size controlled by the voltage applied to the piezo actuator.

**Figure 2.**
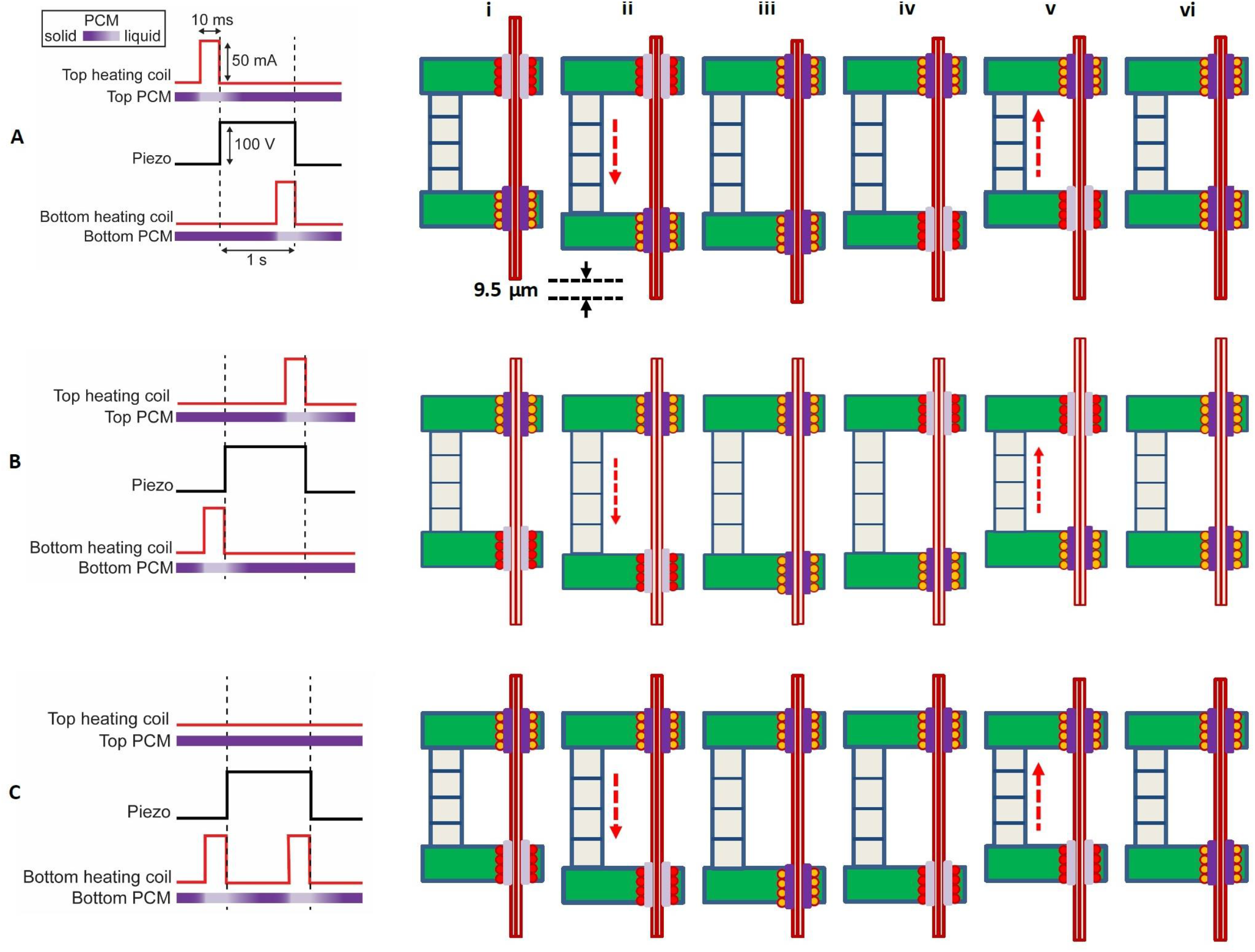
Sequence of electronic actuation steps used by the microdrive for a single translation step. (A) A single downward step (here 9.5 μm) of the probe is performed by six inchworm motor phases (i)-(vi). Long-distance downward probe translation can be accomplished through an unlimited number of repeated single step sequences. In this schematic, the top gripper board is fixed, while the bottom gripper board moves with the piezo extension or contraction. Positive voltage applied to the piezo causes it to extend. (B) A sequence change in the activation of the micro-grippers results in a single upward step. Importantly, the actuation signal to the piezo remains the same. (C) Single no-step sequence, applied to those probes that are to remain in place while other probes are moved in a multi-probe microdrive. The piezo is still actuated, extending and contracting as in (A) and (B), but the sequence of the actuation signals to the top and bottom board heaters is such that the probe does not move. Pulse timing is not shown to scale.

A simple change to the sequence of heater and piezo actuations reverses the translation of the probe. That is, if the bottom micro-gripper is released before the piezo extension, and the top micro-gripper before the contraction, then the motion is reversed, yielding a single upward translation step (Figure 2B). Finally, if only the bottom micro-gripper is released before each piezo actuation, then the probe will be decoupled from the piezo actuation and there will be no translation step of the probe (Figure 2C). The ability to keep probes stationary as the piezo moves is critical to the MPSA microdrive concept, as described in the next section.

### Multi-probe-single-actuator inchworm microdrive operation

The availability of the three distinct probe translation operations of the microdrive while the piezo actuator goes through the same extension/contraction steps (as described in Figure 2) allows the microdrive to independently translate multiple closely-packed, parallel probes while still only using a single piezo actuator. This is accomplished by the placement of multiple micro-grippers into the top and bottom PCBs, such that they are in close proximity and independently electronically controllable for the necessary gripping actuation steps described in Figure 2. The schematic diagram of the MPSA inchworm motor microdrive is shown in Figure 3A, while a photograph of the side view of a model device with three loaded twisted wire tetrode electrodes is shown in Figure 3B. Here, the top gripper board is fixed, while a single piezo actuator moves the bottom gripper board. Independent current sources are connected to each of the helical coil heaters, while hardware and software developed for the microdrive direct the sequence of operations performed by the microdrive’s electronic components (independent top and bottom board helical coil heaters and the piezo actuator). A schematic diagram of the microdrive electronic circuitry is shown in Figure 3C. We have prepared neural robotic microdrives with up to 16 independent helical coil heater micro-grippers in each board, typically in a hexagonal close packed array, with 300μm center-to-center micro-gripper spacing, as shown in Figures 3D-F (see Materials and Methods). It is important to note that the size and the weight of the microdrive remains essentially the same with the increase in the number of loaded neural probes, as the size and weight of additional probes and grippers are nearly negligible. We also note that in addition to the resistive helical coil shown in Figure 1C and 1D, the resistive heater gripper could be potentially constructed from plated or deposited resistive material in the via, a chip resistor installed adjacent to the via, or other manufacturing methods.

**Figure 3.**
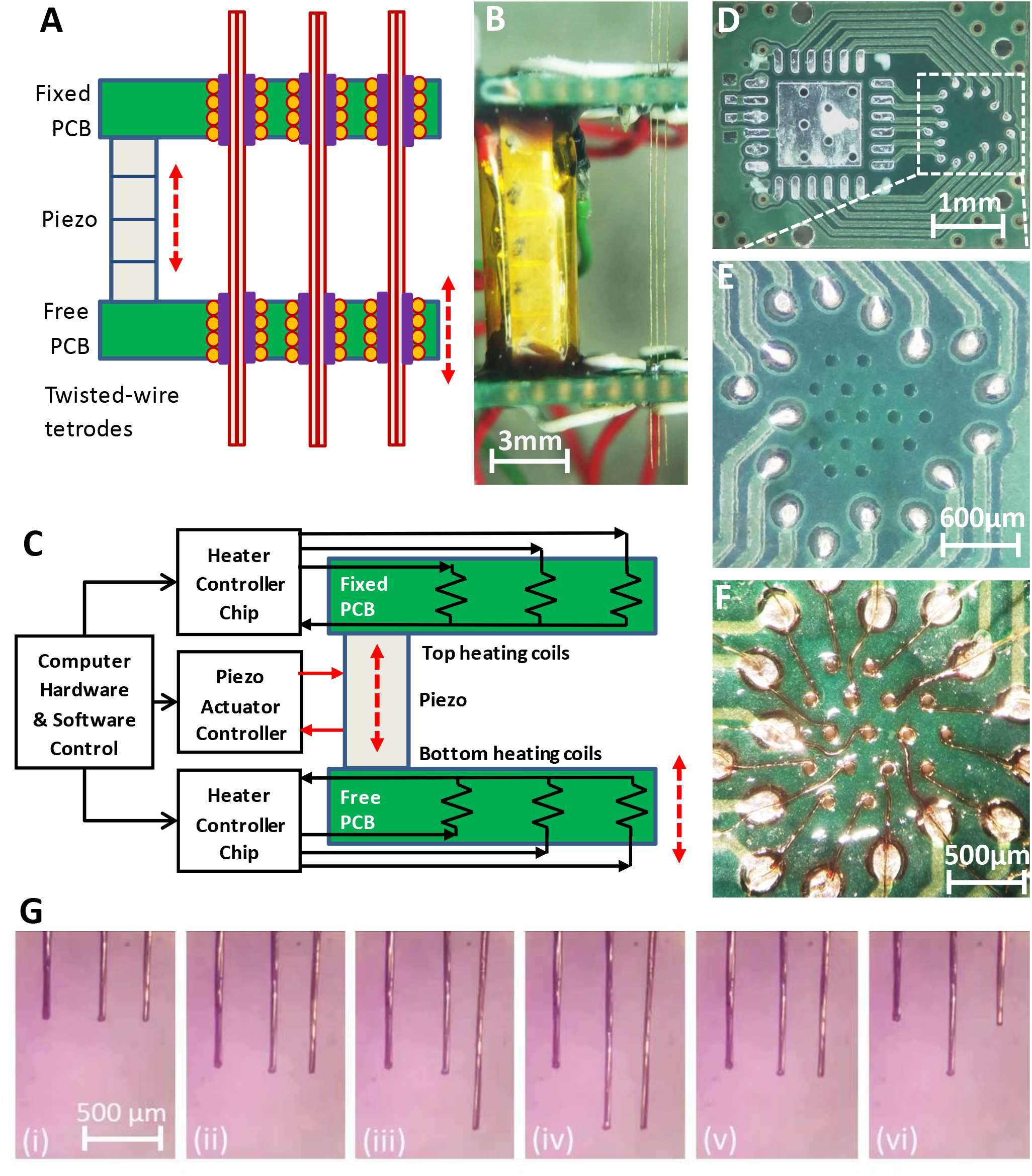
Multi-probe translation capability. (A) An MPSA microdrive with independently movable probes. Axially aligned pairs of independently controlled micro-grippers are integrated into the top and bottom micro-gripper PCBs. The top board is fixed, and the bottom board is free to be moved by the piezo. (B) Side view of a model MPSA microdrive loaded with three tetrodes (right). (C) Schematic diagram of the electronic circuitry. Independent current sources are electrically connected to heater coils in the vias, while hardware and software control the sequence of electronic actuation signals. (D-F) An example unpopulated printed circuit board (D) with 16 drilled vias (E) for placement of 16 micro-grippers in a hexagonal close packed arrangement (F), with 300 μm center-to-center spacing. Demonstration of multiple independent probe movements is shown in Supplementary Video 2 and (G): (i) with all three probes aligned in height, (ii) all three probes moved together downward, (iii) just the right probe is moved downward, while the left and middle are held stationary, (iv) just the middle probe is moved downward, while the left and right are held stationary, (v) the right and middle probes are moved together upward, while the left is held stationary, (vi) the right and left are moved upward, while the middle electrode is held stationary. The piezo actuator always receives the same actuation sequence of signals throughout these different independent electrode translation options (down step, up step, and no step sequences of Figure 2). Forty-two piezo steps were taken between each consecutive image.

In Figure 3G and Supplementary Video 2 we demonstrate the ability of the MPSA microdrive to independently move multiple probes (twisted wire tetrodes). The sequence of signals to the pairs of top and bottom board micro-grippers, as described in Figure 2, independently and simultaneously controls the motion of each probe (Figure 3G).

### Multi-probe-single-actuator inchworm microdrive characterization

To independently move the probes, the micro-grippers must be sufficiently thermally independent. The close packing of probes for many applications, however, could cause significant thermal coupling and risk unintentional probe release. We therefore determined whether and under what conditions the micro-grippers could function in a pattern useful for neural recordings. Simulations and translation testing revealed the importance of the short distance between a micro-gripper’s heater and PCM relative to the distance between micro-grippers (which are separated by PCB laminate, a composite typically having low thermal diffusivity) and that short, controlled heat pulses would be critical (See Materials and Methods). A hexagonal close packed array of 16 tetrode-sized micro-grippers, with 300 μm center-to-center spacing was accomplished when the heating coils (~125 Ω) were operated with ~45 mA, yielding a ~250 mW heat output and release of the docosane-filled micro-grippers in 4 - 12.5 ms depending on their initial temperature. Heating for longer than 12.5 ms from room temperature was functionally unnecessary and risked unintentional release of adjacent grippers. We additionally increased the board cooling and therefore cycle rate by adding internal copper layers to the gripper boards outside of the micro-gripper region. Step rates of 0.5 Hz (~300 μm/min) could be continuously maintained with steady-state board warming of a few degrees and short bursts of up to 2.5 Hz were feasible. Much faster step rates were possible when reduced numbers of probes were used, due to the reduced heat dissipation in the micro-gripper region.

The mechanical characteristics of the MPSA microdrive were quantified with optical metrology (Figure 4A). The probe (tetrode) was moved using the maximum rated voltage for the piezo, producing reliable stepwise motion as expected from the inchworm motor scheme (Figure 4B). Step size can be decreased for finer-resolution motion by adjusting the voltage applied to the piezo during each step. The displacement of the probe matched the actuation distance of the piezo stack throughout the 100 V range of the driving signal (Figure 4C). The minimum reliable step size was measured to be ~600 nm, near the limit of our optical resolving power, and the maximum was 9.5 μm, corresponding to the stroke size of the piezo.

**Figure 4.**
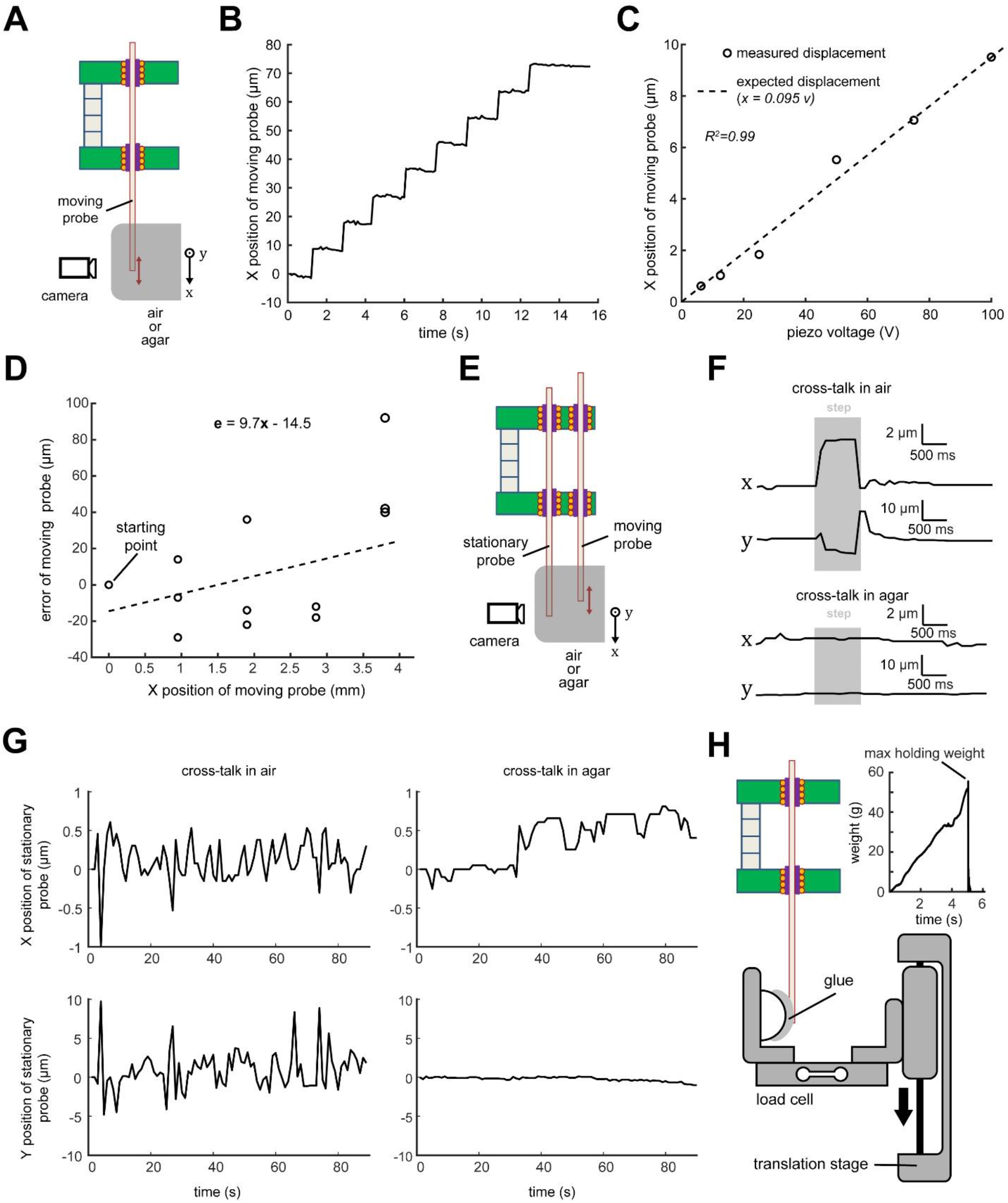
MPSA microdrive mechanical characterization. (A) Experimental setup diagram: microdrive is clamped such that probes move in the field of view of the camera or microscope. In some experiments, probe tips were embedded in agar. (B) Measured stepwise motion of a tetrode loaded into the microdrive. Step size was set to 9.5 μm (maximum). Motion was imaged at 2 frames/s. (C) Correspondence between piezo command voltage and measured tetrode step size. (D) Accuracy of probe motion over 4 mm travel range. (E) Experimental setup for measuring cross-talk between a moving probe and a stationary neighboring probe. (F) Representative displacements in X (on-axis) and Y (off-axis) of a stationary probe during a single step of a neighboring probe. Probes moving in agar brain phantoms (bottom) exhibited smaller displacements than those moving in air (top). Step duration is marked in gray and corresponds to actuation of the piezo stack. (G) Long-term drift of stationary probe during continuous stepping motion of moving probe (9.5 μm step size). (H) Experimental setup for measuring maximum holding weight of a single micro-gripper. A translation stage moved the load cell until the tetrode came loose from the micro-gripper. The maximum holding weight was considered to be the weight supported by the micro-gripper immediately before breaking (inset).

We then characterized the accuracy and repeatability of probe positioning. Accuracy was defined as the difference between a commanded probe displacement (1 mm and 4 mm) and the actual measured probe displacement. Repeatability was defined as the displacement error associated with approaching a single position from different starting locations. A tetrode loaded into the microdrive and advanced by 9.5 μm steps exhibited a mean on-axis accuracy of 7 μm within a 1 mm total displacement range and 58 μm within a 4 mm total displacement range (Figure 4D). Intuitively, this means that if the probe is commanded to move 1 mm, it will overshoot or undershoot by on average of 7 μm. It is expected that accuracy could be increased over a longer travel range with closed loop motion. We subsequently measured repeatability. The tetrode loaded in the microdrive exhibited a mean repeatability of 38.3 μm (range: 5.5 to 106.1 μm, n=6 positions). In an agar brain phantom, the repeatability improved to 4.7 μm (range: 0.9 to 12.9 μm, n=6 positions). A possible cause for the improvement was the reduction of off-axis probe motion in the micro-gripper when the probe tip is constrained in the viscous agar. Intuitively, this means that if the probe is commanded to move to a specific position many times, the spread of the resulting probe positions will be on average 4.7 μm in agar.

Any coupling in the motion between adjacent probes is undesirable; we therefore measured the effect of moving one tetrode on a neighboring stationary tetrode positioned 300 μm away (Figure 4E). When both tetrodes are in air, a single 9.5 μm step of the moving tetrode caused the stationary one to move momentarily 3.6 ± 2.1 μm on-axis and 23.8 ± 4.4 μm (n=5 steps) off-axis. We reasoned that these momentary deflections of the stationary tetrode are likely caused by a combination of (1) thermal and phase change expansion of the PCM within the released bottom micro-gripper, (2) lateral motion within the bore while released and (3) the non-straight and non-cylindrical tetrode sliding against the inner wall of this micro-gripper. We therefore hypothesized that much of this cross-talk could be reduced by mechanically dampening the tetrode tip, as would occur in the brain. As expected, when the same test was performed in an agar brain phantom, the stationary tetrode moved significantly less (x: 1.1 ± 0.2 μm, p=0.03; y: 0.85 ± 0.2 μm, p<0.0001; n=5 steps; Student’s unpaired t test; Figure 4F, Supplementary Video 3). In a separate experiment, we continuously moved a tetrode and quantified the slow drift of the neighboring stationary tetrode. After the moving tetrode was stepped continuously for 90 s (total distance of 500 μm), the stationary one was found to have drifted less than 2 μm in X and Y in air as well as in agar (Figure 4G).

Finally, the maximum weight that can be supported by the microdrive grip on a tetrode before the micro-gripper hold breaks was measured to be 58.0 ± 2.4 g (n=5 tetrodes; Figure 4H), equivalent to a holding force of 0.57 N.

### Multi-probe-single-actuator inchworm microdrive neural recordings

An integrated robotic microdrive for neural recordings with 16 twisted wire tetrode electrodes was assembled as shown schematically in Figure 5A, and in the photograph of Figure 5B. In addition to the main MPSA inchworm microdrive structure described previously, the complete design included an electrode interface and control board (EICB) with 64 recording channels (4 channels per each tetrode), connectors for extracellular recording amplifiers, electrode cannulas for maintaining electrode alignment, and a mechanical cage and mounts for encasing and positioning the drive for acute and chronic recordings. We again demonstrated the independent translation capability of the microdrive using only a single piezo actuator with 16 twisted wire tetrodes (Figure 5C, Supplementary Video 4). The sequence of electronic actuation signals sent to the sets of 16 micro-grippers in the top and bottom boards (as described in Figure 2) independently controlled the direction of motion of each twisted wire tetrode.

**Figure 5.**
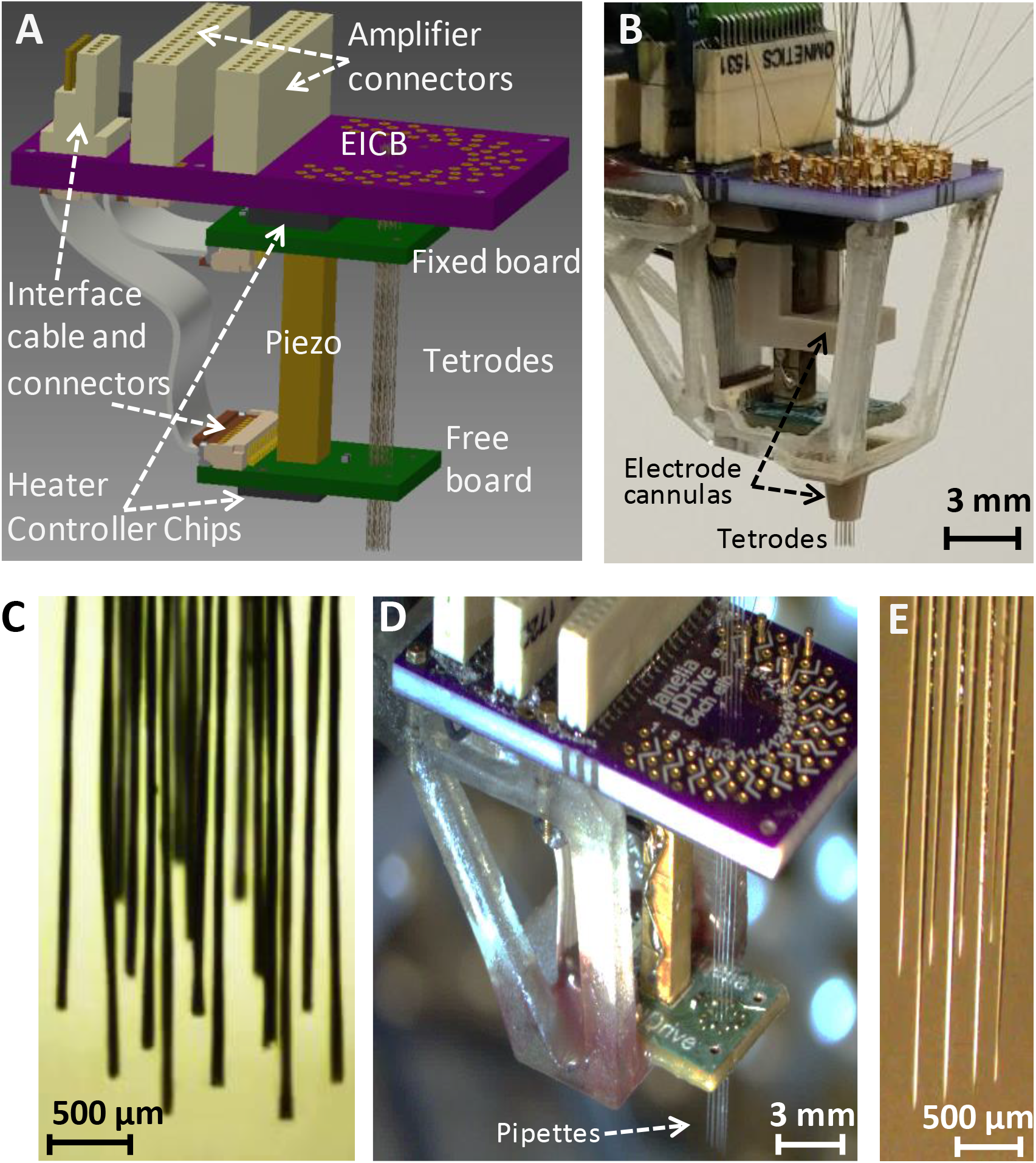
Implantable MPSA microdrives for neural recordings. (A) Schematic showing the main MPSA microdrive with the added EICB. (B) Photograph of the implantable tetrode-drive for acute neural recordings. Cannulas were added to increase tetrode stiffness during insertion. A 3D-printed cage was added on the exterior for protection and ease of handling. The tetrode-drive was used to independently position 16 independent tetrodes shown in (C) using a single piezo actuator (Supplementary Video 4). (D) Pipette-drive loaded with 8 glass micropipettes, and independent motion of each micropipette (E) was also demonstrated (Supplementary Video 5).

As mentioned earlier, the micro-grippers of the device can grip probes of any material and cross-sectional shape, as the liquid PCM in the gripper bore conforms to the shape of the probe before solidifying. For the purposes of demonstrating this feature of the microdrive, in addition to neural twisted wire tetrode electrodes, we also independently translated 8 glass micropipettes (50 μm inner diameter / 80 μm outer diameter) while only modifying the size of the heater coil (~100 μm inner diameter / 135 μm outer diameter) (Figure 5D-E, Supplementary Video 5).

We first demonstrated the neural recording capability of our microdrive in acute surgical settings on anesthetized rats (see Materials and Methods), initially with 4 twisted wire tetrodes (Supplementary Video 6), and then with 16 twisted wire tetrodes as shown in Figure 6A. Under remote operation, the electrodes (Figure 6B) were independently advanced in 9.5 μm steps sequentially into the brain, while 64 channels (from 16 tetrodes) of neural signals were monitored visually in order to guide the approach of the tetrodes towards the targeted CA1 layer of the rat hippocampus. Figure 6C shows the sequential tuning of 16 tetrodes to the CA1 layer of the rat hippocampus. In the first sub-panel, the 16 twisted wire tetrodes begin just above the CA1 layer, where they detect mainly the local field potential fluctuation in the brain and only small neural spikes. The sub-panels that follow show clearly distinguishable spiking activity after the tuning of 7, 12, and all 16 tetrodes. Importantly, tuning of additional tetrodes did not compromise the quality of the already placed tetrodes, as desired. In the fourth panel, the recordings after all 16 twisted wire tetrodes had been tuned is shown, with high-quality neural spike signals from all 64 channels.

**Figure 6.**
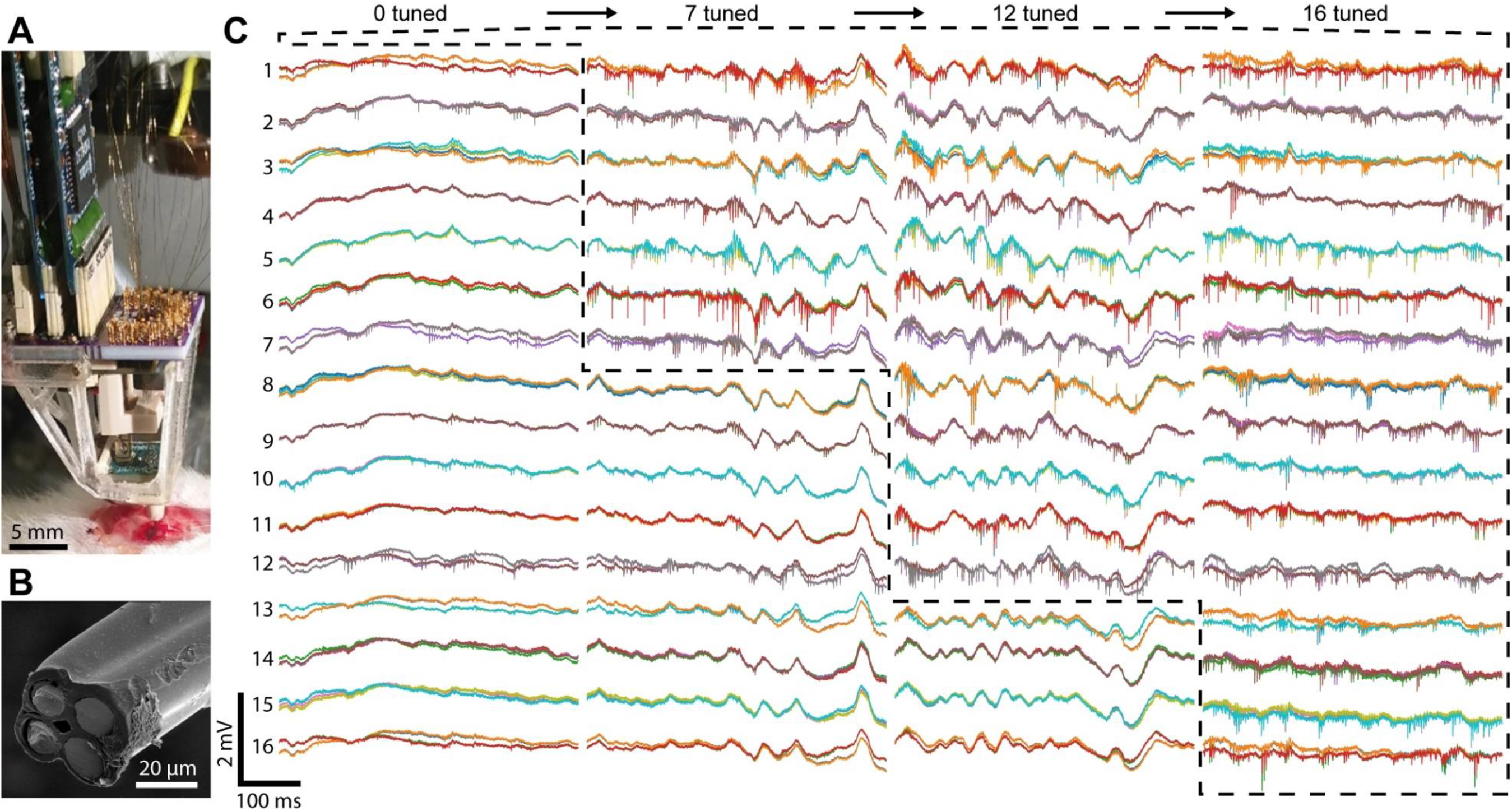
Neural recording with the 16-channel implantable tetrode-drive in an acute surgical setting. (A) Photograph of the acute experiment (craniotomy at bottom). (B) Scanning electron microscope image of the cut-face of a tetrode with the 4 exposed neural recording sites. The tetrodes were independently advanced in 9.5 μm steps sequentially into the brain. (C) Voltage traces during precise depth tuning, beginning after all tetrodes were brought near CA1. The position of each tetrode was independently adjusted to optimize its signal; dashed lines show groups of probes that have been optimally tuned. Previously tuned tetrodes did not lose signal after tuning of neighboring electrodes. For each of the 16 tetrodes, the 4 traces from each electrode (different color for each) of a tetrode are overlaid.

As a final demonstration we performed chronic neural recordings in awake rats using an implantable tetrode-drive with 4 loaded tetrodes (Figure 7A). Before application of dental acrylic to cement the device to the skull, the overall weight was 4.5 g with overall dimensions of 25 × 15 × 31 mm. As mentioned above, the weight and size would not have changed significantly if all 16 probes had been loaded. The microdrive was maintained for 6 weeks and the tetrode position was periodically adjusted through remote electronic control for fine tuning of neural spike signal quality. The recordings exhibited expected sharp-wave ripple and spiking activity (Figure 7C-D).

**Figure 7.**
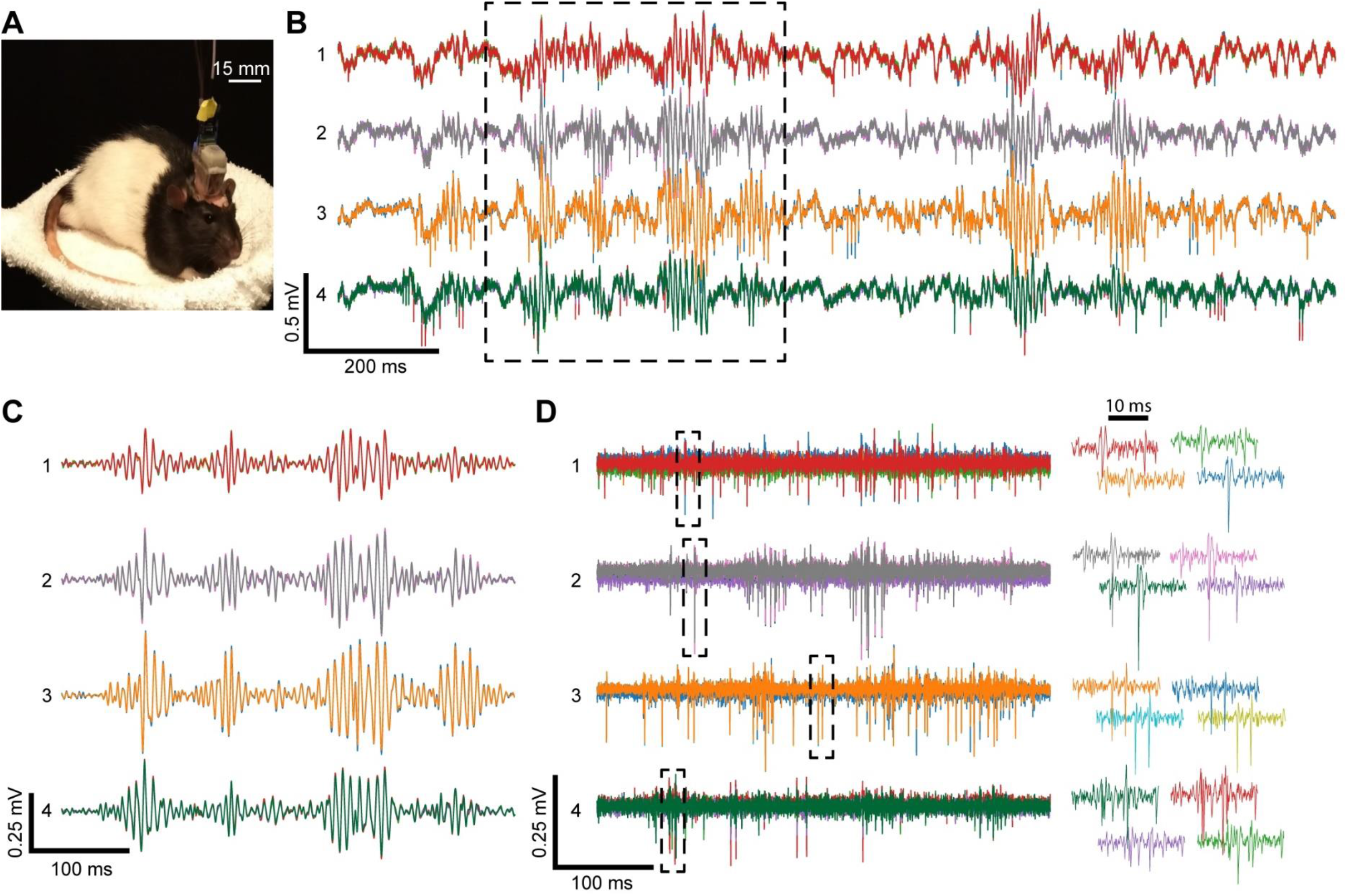
Chronically implantable head-mounted MPSA tetrode-drive. (A) Rat implanted with the tetrode-drive. (B) Sample raw voltage traces from 4 tetrodes obtained ~6 weeks after implantation showing LFP, sharp wave ripples and spiking activity in the hippocampus. (C) Boxed voltage segment from (B) filtered to isolate sharp-wave ripples (bandpass: 100-300 Hz). (D) Boxed voltage segment from (B) filtered to isolate action potentials (bandpass: 500-8000 Hz). For each of the 4 tetrodes, the 4 traces from each electrode (different color for each) of a tetrode are overlaid. Right: spike waveforms from boxes at left.

## Discussion

We described the concept and use of a new robotic MPSA neural microdrive. The microdrive features a novel electronically-controlled micro-gripper that utilizes temperature controlled phase change material as a gripping medium^61^. Using the gripper’s microscopic dimensions, independent electronic actuation control, and high packing density, we demonstrated robotic placement of multiple independent neural recording electrodes into the CA1 region of the rat hippocampus. This was done with micrometer precision with the unlimited probe translation capability, using only a single piezo actuator in acute and chronic *in vivo* settings.

We highlight several advantages of our MPSA concept and implementation over traditional neural microdrives. First, the travel range of the MPSA microdrive is conceptually only limited by the length of the probe. Therefore, the MPSA microdrive could be used to simultaneously target distant regions of the brain (for example, cortical and thalamic brain tissue) in large rodents or even primates^64^. Second, the MPSA concept makes the microdrive inherently more scalable than traditional microdrives. Each additional channel in a traditional motorized microdrive typically requires a motor, shuttle, and connector, causing the weight and size to increase substantially with channel count. On the other hand, in the MPSA microdrive, the addition of an extra channel causes a negligible weight and size increase, since the need for independent actuators is removed. Third, further improvements in scalability will rely on advances in the miniaturization of microchips and other electronic components (e.g. higher-channel-count current source integrated circuits), which is likely to outpace the miniaturization of traditional motors. Fourth, traditional microdrives utilize an inverted cone geometry in order to accommodate bulky actuators. This arrangement accommodates the use of flexible neural probes such as twisted-wire tetrodes and long silicon shanks^65^, but precludes the use of stiff micropipettes and short silicon shanks which are typically too inflexible to bend along the angular guide channel and through the vertical cannula. Since the micro-grippers in the MPSA microdrive can be tightly packed, the conical geometry is not needed and rigid probes such as micropipettes can be translated.

Minimal thermal design is required for a single probe implementation but coupling and heat flow become more important as the probe count and heater density increases. The relevant phenomena, however, can be readily modeled with heat transfer simulations and we envision that more sophisticated mechanical and control schemes could yield higher probe densities and movement speeds. In particular, increasing the movement speed is important for generalizing the use of these temperature-sensitive PCM based grippers, as many applications require faster rates than utilized here. Examples of more advanced methods include integrating copper heat sink regions between micro-grippers and optimizing heating timings based on the mutual impulse response and heating history of all the thermally coupled heaters on a board. Additionally, the PCM could be chosen for the temperature of the application or designed to have properties helpful to the micro-gripper functionality such as higher thermal diffusivity and a narrower phase transition^66^. While this MPSA microdrive design is particularly attractive for rodent electrophysiology where miniaturization is critical, in other uses, weight and size can be traded for additional functionality. For example, in situations where motion accuracy is critical, a positional feedback system such as a small camera could be installed. Alternatively, the design does not require ferrous components, and therefore an MRI-compatible version of the microdrive^67^ would be a particularly interesting variant of imaging feedback control. Further directions for the implantable neural MPSA microdrive include enabling untethered control and recordings through wireless communication via Bluetooth or Wi-Fi, implementing on-board piezoelectric drivers, and battery-powered operation.

An important advancement to the functionality would be the implementation of autonomous probe positioning. For example, closed-loop spike locating and fidelity monitoring capabilities would allow for the placement of a probe within a region of interest based upon a model signal and thereafter optimization of signal quality, in which degradation could trigger automatic compensatory motion^51^. Such automation could lead to both improved data yield and reduced operator time in experiments.

Apart from neural electrophysiology probes, the MPSA microdrive could also be used for other biological as well as non-biological probe translation tasks^67,68^. For example, the micro-grippers could be sized to accommodate hypodermic needles, optical fibers and ultrasonic transducers and sensors of a variety of scales. In conclusion, our robotic microdrive provides the capability for independent positioning of multiple parallel probes using only a single piezo actuator with significant benefits wherever the need arises for small weight, miniature device size, sub-micron level probe positioning precision, and remote control.

## Materials and methods

### Micro-gripper construction

Heating coils were prepared by winding fine insulated nichrome wire (California Fine Wire, Model Stableohm 800A, 12.5 μm diameter conductor, 17.5 μm diameter with insulation) around a 75 μm diameter Tungsten wire core (California Fine Wire). The final micro-heater had ~45 turns, as shown in Figure 1C, and had a typical resistance of 125 Ω for a tetrode micro-gripper. Docosane (Sigma-Aldrich, item 134457) was chosen as the PCM due to its non-toxicity and sharp, low melting temperature (~42 °C), which is still above animal body temperature^69^.

### Thermal characterization

The gripper board designs were simulated in COMSOL Multiphysics before fabrication in order to implement a design with sufficient micro-gripper thermal independence (See Supplementary Information). Independence was assessed based on whether the temperature in the ‘off’ micro - grippers remained below the lowest temperature of the phase transition region (~41 °C) when the ‘on’ micro-grippers could reach a temperature above the highest temperature of the phase transition region (~44.5 °C). An additional safety margin of ~5 °C for both temperatures was included when analyzing the simulations to conservatively account for potential differences between the model and the fabricated device. Simulations were performed with an initial temperature of 20 °C, and a smaller safety margin would be used for initial temperatures closer to the phase transition temperature to reflect the smaller overall changes. Heat duration profiles as a function of board temperature were also obtained due to expected temperature changes in the environment and during use. Timing parameters were ultimately adjusted based upon empirical movement tests with assembled microdrives of varying probe counts.

### MPSA microdrive motor assembly

Gripper boards were manufactured using standard PCB technology (8 × 11 × 0.8 mm, 6 layers, Sierra Circuits). Each board contained an integrated current source chip (STMicroelectronics LED1642GW), associated passives, 0201 thermistor, and a flat flex cable connector. Each current source chip was pre-characterized in a custom, automated setup in order to use only chips with consistent and high current across all channels. These components were soldered onto the board before drilling of the micro-gripper vias. To increase the micro-gripper and overall board cooling rate (and therefore the cycle rate), solid internal layers and thermal vias (~27 per cm^2^) in the non-gripper regions of the board were used to increase the thermal conduction across the plane and through the thickness of the board, respectively (see Supplementary Information). Additional through-holes were placed on the board and used for alignment and mounting purposes when integrating these boards into larger structures. Via holes (4.7 mil for tetrodes, 5.9 mil for pipettes) were drilled into the PCB in a hexagonal close packed array with 300 μm spacing using a 5-axis CNC mill (Hermle). The micro-heaters were inserted into the via, glued in place with high temperature epoxy (Thorlabs, 353NDPK), and the core Tungsten wire was extracted for a final open bore resistive via, shown in Figure 1D. The heating coils in each board were individually soldered to exposed pads which were routed to the independent channels of the current source chip. The connection pads and leads were insulated with epoxy in order to yield a device which was mechanically robust to handling and contact near the heaters, as would occur during the loading and cleaning procedures. To complete the microdrive assembly, the piezoelectric stack (Thorlabs PK3CMP2) was positioned on the designated locations on each board and epoxied to each, one at a time. A custom jig with alignment pins (McMaster-Carr) was used during the curing process in order to ensure that the boards were aligned and parallel.

### Implantable microdrive system

In addition to the MPSA motor described above, the implantable microdrives (tetrode-drive and pipette-drive) necessitated an EICB, a base station, and a PC application for control.

The EICB provided crimping locations for 64 electrodes and 2 Omnetics connectors for 32 channel Intan amplifier boards (Intan Technologies, C3314, Figure 5B). These were ordered in 16 sets of 4, for a maximum of 16 tetrodes. There were additional contact locations for reference electrodes and grounding connections. In the center of the connection region a pair of alignment holes were included to match the ones used on the gripper board. This facilitated alignment of the boards and the drilling of probe vias in the EICB.

The control section of the board consisted of a microcontroller (Microchip PIC16F18345), 3-color LED indicator (Rohm Semiconductor SMLP36RGB2W3R), and connectors for the base station cable as well as the two flex cables going to the heater boards and connection pads for the piezo wires. The microcontroller implemented the timing and coil heating operations determined by the movement set in the PC software. Thermistors were used to measure the temperature of each board to account for onboard heat dissipation and compensate for animal body heat. The temperature was fed back to an algorithm in the PC software and used to adjust heat durations. The cable to the base station consisted of 5 meters or shorter lengths of thin, unbundled wires (Cooner Wire, Stranded Bare Copper FEP Hookup Wire).For connecting the gripper boards to the EICB, custom length flat-flex cables were made by shortening a silicone flex cable to the proper length, and adhering plastic backing to allow insertion into the connectors.

The base station consisted of a USB connected microcontroller (Arduino Mega 2560), piezo driver (Thorlabs KPZ101), heater power circuit and connectors. A trigger line from the EICB was monitored by the base station microcontroller in order to command the piezo driver with highly accurate timing.

The PC control application was implemented in LabVIEW (National Instruments). It provided the ability to select the probes to move and the direction to move each probe. The application had the capability for scripted series of movements, changing the frequency of movements, the current through the heating coils, durations of heating, and temperature compensation. It also produced a record of the movements and displayed the total movements and offset from a set point.

### Implantable microdrive assembly

For the tetrode-drive, a custom-milled, temperature stable polyether ether ketone (PEEK) spacer was used to join the top gripper board and EICB to a 3D printed superstructure cage, which protected the microdrive from damage and allowed for easier handling. Although the high aspect ratio of the bores maintains good probe axial alignment in the micro-grippers, we realized that flexible nichrome wire tetrodes bend easily, which can reduce or negate the length of each step. We therefore included two guiding PEEK cannulas. The mid-drive and bottom-tip cannulas were aligned with the top and bottom micro-gripper vias using guide pins and attached to the top board and cage, respectively. The bottom board was chosen to be mobile in order to minimize bending by minimizing the length of tetrode between the piezo-coupled micro-gripper and the brain. For chronic testing, the EICB was also enclosed with 3D printed caps, and the open sides of the cage were sealed with plastic film sheets. The mid-drive cannula was not used in the chronic experiment. M0.6 fasteners (Prime-Miniatures) were used throughout for modifiable attachments.

During acute experiments, it was seen that evaporation from the craniotomy condensed on the bottom board. For this reason, the bottom board was insulated with a conformal silicone PCB coating (MG Chemicals 422B) throughout the non-heating area.

### Probe loading

Twisted wire tetrode recording electrodes were prepared from fine insulated wires (California Fine Wire, Model Stableohm 800A, 12.5 μm diameter conductor, 17.5 μm diameter with insulation). Each electrode was loaded into the microdrive by threading it through the empty micro-heater bores and cannulas with fine tweezers under an inspection microscope. A small particle of solid PCM (sized to underfill the probe filled bore) was placed near the micro-gripper bore either on the board or on the threaded probe with fine tweezers. The PCM was melted either using the micro-gripper’s heater or a heat lamp so that it could flow into and fill the bore through capillary action. One end of the tetrode was cut with a sharp razor blade in order to expose the 4 neural recording sites (shown in Figure 6B), while on the other end each wire was crimped separately into the EICB for connection to the digital recording system (Figure 5B). When loading many probes at high density, the probes were left loose in their bores until all were threaded, after which the PCM on the probes was melted and slid into the bores. Thereafter, the PCM in the micro-grippers was melted during any additional probe handling (e.g. trimming and site exposure). Previously leaving the probes mobile in the bores avoided damage to the probes due to contact with the tweezers and also better controlled the amount of PCM in each micro-gripper by preventing flow into neighboring bores. Because the EICB did not need to be enclosed during acute experiments, the tetrodes were left long between the crimp locations and the EICB probe vias.

The same procedure was used to load the microdrive with the glass micro-capillaries (Vitrocom Model CV0508 50 μm ID / 80 μm OD borosilicate glass) for the glass capillary microdrive translation demonstration.

### Mechanical testing setup

To characterize the motion of the microdrive, it was first rigidly mounted in a 3D-printed clamp such that probes moved horizontally. Probes (pipettes or tetrodes) were loaded into the drive as described above. The maximum piezo displacement voltage (100 V) was used for all tests unless otherwise noted. For accuracy characterization, the MPSA microdrive loaded with a single tetrode was placed under a USB microscope (VMS-004, Veho), such that the probe was in the field of view. In all other motion characterization experiments (repeatability, resolution, step size, cross-talk), the assembly was placed under a probe station microscope (Alessi REL-4100A) equipped with 10X and 60X objectives (Mitutoyo) and a digital camera (EO USB 2.0, Edmund Optics). In the cross-talk experiments, the microdrive was loaded with two tetrodes. In a subset of experiments, probes moved through 0.2% w/v agarose (Millipore Sigma). Bidirectional repeatability was measured by repeatedly commanding a probe to move to a single target position in the travel range. The target position was approached from 0.4 to 5 mm away from both directions.

For the micro-gripper holding force characterization, we used a load cell (RB-Phi-203, RobotShop), which we first calibrated with known weights. One end of the load cell was attached perpendicular to a linear translation stage (PT1, ThorLabs). A 3D-printed bracket with a metal bolt was attached to the other end. The free end of a tetrode in a stationary microdrive was superglued to the bolt. The translation stage was used to pull the tetrode until it came loose from the micro-gripper. The maximum force on the load cell immediately before the break was considered to be the holding force of the microdrive. All image analysis was performed using FIJI^70^.

### In vivo electrophysiology

All procedures involving animals were performed according to methods approved by the Janelia Institutional Animal Care and Use Committee. All acute surgical experiments were performed on anaesthetized Wistar male rats (age: P20-P30). The animals were placed in a stereotaxic apparatus maintained at 37°C with the anesthesia maintained at 1.5-2% isoflurane. A small craniotomy (diameter ~1mm) was drilled in the skull (position coordinates for dorsal hippocampus: 3.5mm posterior of bregma, 2.5mm lateral of midline) and the dura removed with a sharp needle in a small region (diameter ~0.25mm) centered in the craniotomy. A chlorided silver wire was folded under a flap of skin near the incision and used as a reference electrode for the recordings. The microdrive loaded with 4 or 16 tetrodes was positioned over the center of the craniotomy using a motorized micromanipulator (Luigs & Neumann) and a 3D printed holder.

For chronic recordings, Long-Evans rats were first anesthetized with isoflurane and headfixed in a stereotaxic frame. A craniotomy was made over the CA1 field of the right dorsal hippocampus (AP −3.8 mm, ML 2.4 mm) and the dura was removed. The tetrode tips were previously gold-plated to reduce the impedance of each channel to < 250 kΩ at 1 kHz, and a stainless steel screw placed in contact with the left cerebellum served as the reference electrode for recordings. Before implantation, the tetrode tips were retracted 50 μm into the bottom cannula of the microdrive which was then filled with melted Vaseline to prevent clogging by blood or CSF. In order to prevent Vaseline from contacting the micro-grippers, the retraction of the probes was limited to 2.5 mm from their deepest point. The bottom cannula was then aligned to the craniotomy using a stereotaxic holder which connected to one of the Omnetics amplifier connectors of the EICB, and the microdrive cage was fixed on the skull with OptiBond (Kerr), A1 Charisma (Kulzer) and dental cement (Lang). Soon after implantation, the depth of sharp wave ripples was found for each probe, which were then retracted and then re-descended over several weeks until sharp-wave ripples and spikes were clearly seen^71^. At the termination point of a chronic experiment, the EICB and microdrive motor were recovered for future use.

Neural recording data from 64 total channels from 16 tetrodes was digitally sampled at rates of 20 kHz/channel for acute experiments and streamed to the computer via USB using an Intan USB Interface Board. For chronic experiments, 16 total channels from 4 tetrodes were digitally sampled at 30 kHz/channel and streamed to the computer via USB using an OpenEphys Acquisition Board.

## Acknowledgments

This work was supported by the Howard Hughes Medical Institute. We would like thank Bill Biddle for excellent scientific machining of the robotic microdrive parts, Brian Barbarits for help with extracellular electrode recording electronics, Steve Sawtelle for help with electronics and PCB design, Roger Rogers for help with 3D printing, and Jeff Magee and David Hunt for helpful discussions.

## Author contributions

M.B. concieved the principal ideas of the phase-change-material heat activated via micro-gripper and multi-probe-single-actuator inchworm microdrive. R.D.S. performed the thermal design and testing of the resistive heating coil micro-grippers and PCM, performed the circuit, microcontroller and PCB design of the gripper boards, EICBs and base station, designed the mechanical and system structure of the microdrive, and developed the software control for remotely operating the robotic microdrive. M.B. fabricated and installed helical micro-gripper heaters. R.D.S. connected the micro-grippers, and assembled all the microdrives. R.D.S. loaded the microdrives with neural recording twisted wire tetrodes and glass micro-capillaries. I.K. performed the micro-gripper and microdrive mechanical and probe positioning, coupling, and gripping characterization. S.T. prepared the twisted wire terodes neural electrodes. M.B. and R.D.S. performed the acute surgeries and neural recordings experiments. S. T. and R.D.S. performed the chronic surgeries and neural recordings experiments. R.D.S., I.K. and S.T. performed data analysis and data visualization. A.K.L and T.H. supervised the project and advised on the surgical procedures and neural recordings. M.B. wrote the initial draft of the manuscript. M.B., R.D.S and I.K. expanded and edited the manuscript with input from all authors.

## Supplementary Materials

Videos: single tetrode, three tetrodes in air, three tetrodes in agar,16 tetrodes, 8 pipettes, 4 tetrodes rat brain insertion through craniotomy, gripper board 15-on-1-off thermal simulation, center cut-line of 15-on-1-off thermal simulation.

## Supplementary Information

### Gripper Board Inner Layers and Coil Via Gap

The gripper board contained 4 solid inner layers of copper for distributing the heat after each pulse with a gap around the via pattern of 0.25 mm (Figure S1). Due to the thermal diffusion time through the PCB laminate, the added copper at this distance would not practically affect the dynamics of the millisecond scale heating process, but would contribute to cooling the gripper region over the one hundred millisecond and greater time scale applicable to continuous stepping.

**Figure S1.**
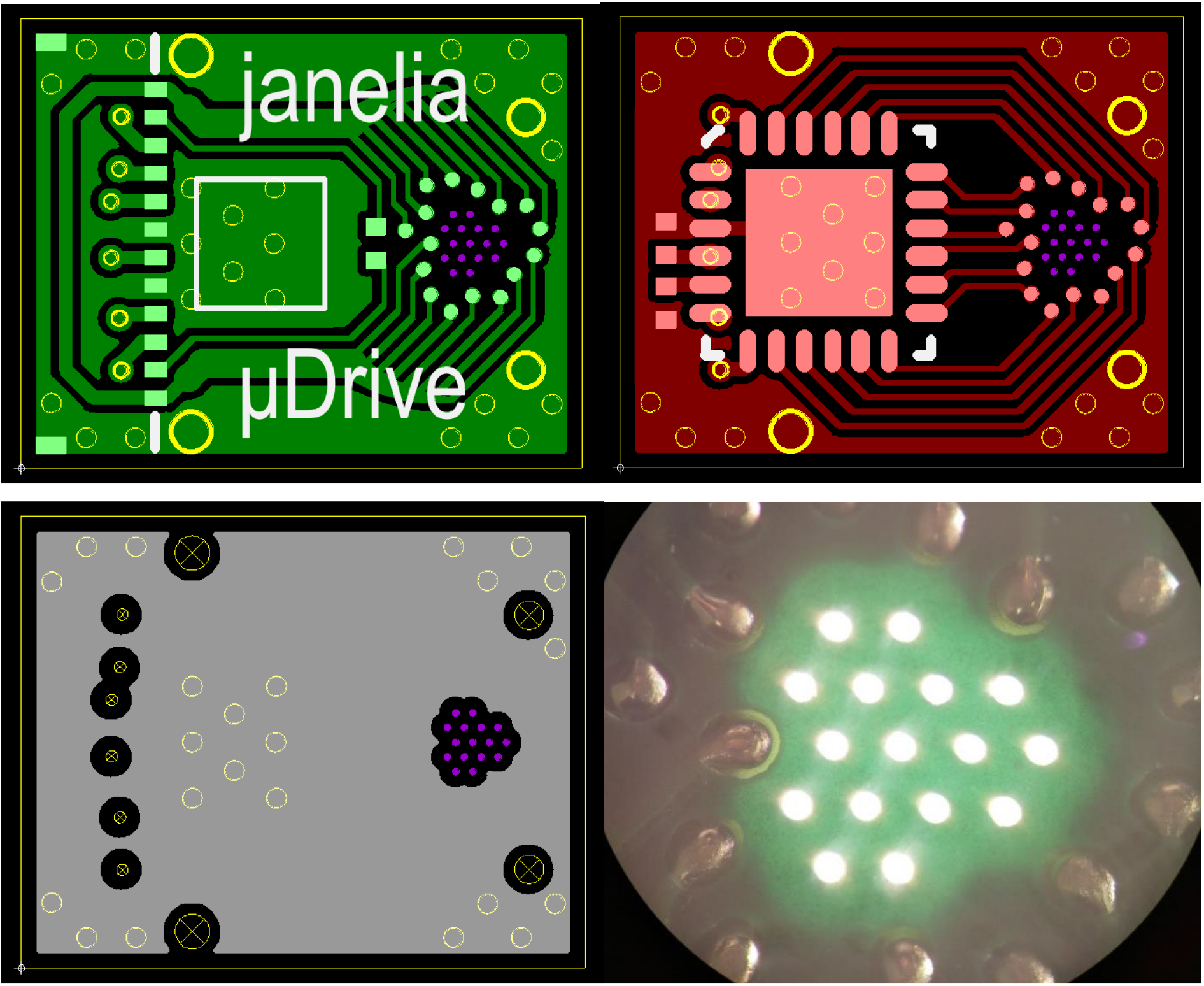
CAD images for the (Top Left) top, (Top Right) bottom and (Bottom Left) inner layers of the gripper board, as well as (Bottom Right) an image of light passing through drilled vias and the non-copper areas of a fabricated gripper board

### Coil Installation Details

Before insertion of the coils, trenches for the leads between the vias and their connection pads were laser-cut into the top surface of the PCB soldermask (Figure S2). After removal of the tungsten pin, the wire leads were placed into the trenches and covered with epoxy. This prevented damage to the fragile coil leads and outflow of PCM along the lead during heating. The insulation of the leads at the pads was then removed using the laser cutter. Since the coil wire metal does not wet solder well, the bare section of the leads was instead forced into a melted bead of solder on the pad, which would solidify and yield a compact electrical connection of negligible resistance. A controlled amount of solder on each pad was obtained from 0.25 mm diameter solder balls (Chip Quik Inc.), and a narrow tip soldering iron was used to prevent bridging between adjacent pads. After soldering, the boards were sonicated in isopropyl alcohol to remove flux residue.

**Figure S2.**
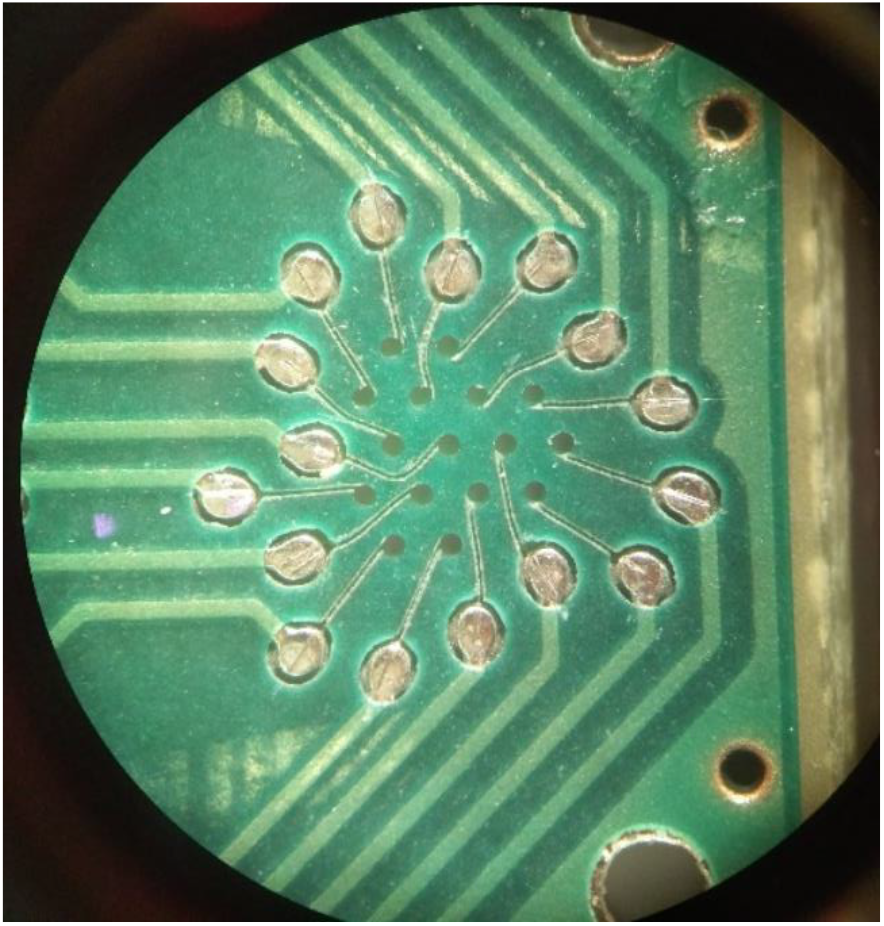
Image depicting the trenches cut into the PCB solder mask for the heater coil leads

### Example Timing of a Step Sequence

For an implanted microdrive where the lower board nearer the animal is typically warmer than the upper board, the heating period for the lower board would be shorter, but both boards would end heating at the same time. For example, based on the board temperatures measured at their thermistor, the lower board may require 8 ms of heating and the top board 10 ms. In order to end at the same time, the lower board would begin heating 2 ms after the top board. Due to the propagation delay of heat through the PCM, a 2 ms delay was applied after the ending of heating before the piezo was actuated. For a 1 Hz step rate, the next heating would begin 0.5 s after the first.

### Pipette-drive Structure Difference

As stated in the main text, the choice of moving and stationary gripper boards is dictated by the application. For the pipette-drive, we wanted to ensure that pipette tips would remain as stable as possible for potential juxtacellular or intracellular electrophysiology experiments. Therefore, in that design, the top board was mobile, while the bottom gripper board was fixed to a 3D printed superstructure which also held the EICB. In this configuration, the nearest board to the tips never releases a non-translating probe, whereas in the tetrode drive, the gripper in the lower board would heat and slide along the probe. No cannulas were added since the pipettes did not bow in practice.

### Potential Failure modes

We note two potential failure modes which become more likely with longer movement lengths: (1) PCM may leak out from the micro-grippers during use and (2) any liquid or tissue drawn up from the brain into a micro-gripper could render it inoperable. Both of these failure modes would be rare and preventable.

Leaking can be prevented by ensuring that the bore was not overfilled, and a sufficient amount of heat was applied to melt all of the PCM during the release. It is important to note that a portion of PCM leaking out of the bore does not prevent the gripper from continuing to function. Additionally, if the PCM that leaked out onto the probe was still near the board, it could be withdrawn back into the bore. This would be done by turning on all the heaters of one board at a low intensity such that the bores and probes would rise above the melting temperature, which would cause the PCM to flow back into the bore. Since the heaters of the other board could remain off, the probes would remain in-place such that this could be done post implantation. The heating and cooling times of this process would be substantial (~10 s) but could be integrated into an overall movement scheme if PCM loss mitigation was critical.

Prevention of foreign substances from reaching the heaters was accomplished by maintaining a maximum total retraction distance from the maximum depth at any point. During chronic implantation, this was done to prevent Vaseline from contacting the PCM, which it could dissolve or at least broaden the phase change temperature profile. Analogous limits could be set when doing acute implantations with or without the tip cannula.

### Gripper Board Simulation

The gripper board designs were simulated before fabrication in order to implement a design with sufficient micro-gripper thermal independence. The simulations were performed in COMSOL Multiphysics using the Heat Transfer module with the phase change material function. Because the region of interest minimally varies through the thickness of the board, 2D simulations were used and set at the plane of an inner layer (Figure S3). The coils were modeled as ring-shaped heat sources, and their heat generation properties were applied as a volumetric average from their measured resistance and the currents passed. 250 mW was typically used, as it is the maximum heat output for the combination of coil and current source used. Lower heat rates corresponded to reduced independence due to heat diffusing out of the gripper. A 3 mil inner diameter was used for the bore, and a circular approximation of a tetrode was used with a 39 μm diameter. Polyimide and epoxy layers were added based on the coil, wire and via dimensions.

**Figure S3.**
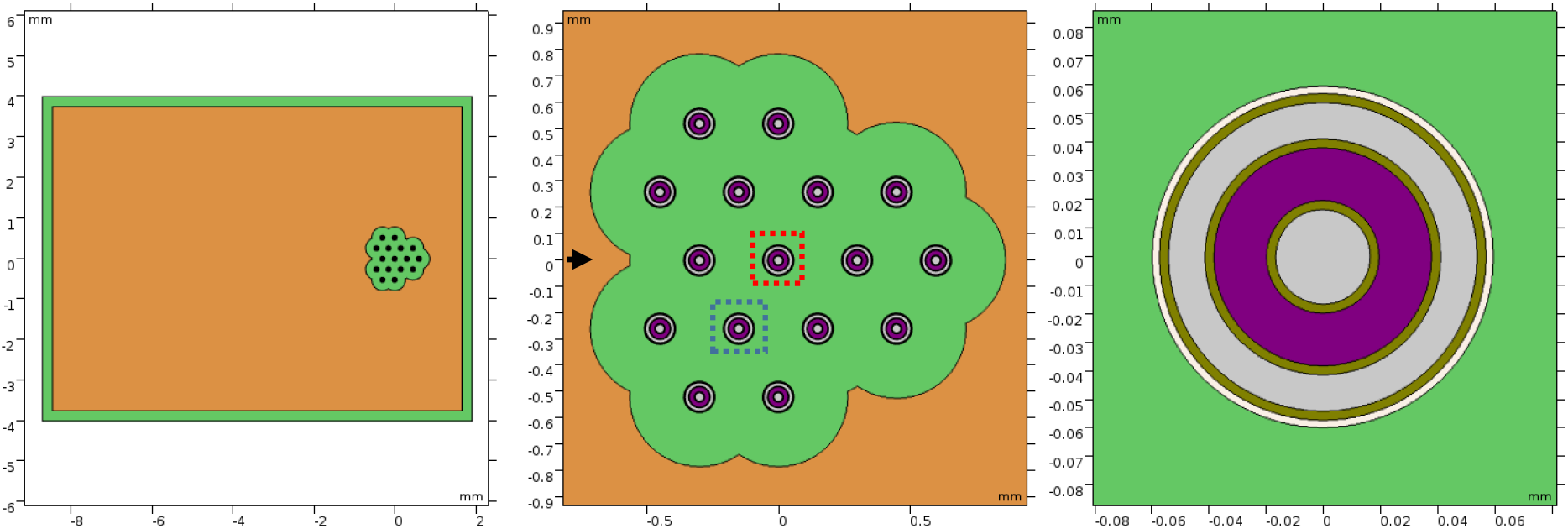
Simulation model geometry. (A) Gripper board. (B) Gripper region with the plotted central gripper and adjacent gripper boxed in red and blue respectively and 1D data cutline axis indicated. (C) Gripper model.

The material properties were implemented using the material option for copper, as well custom materials for the FR-4, epoxy, coil heaters, PCM and probes. In particular, the coil heaters and tetrode probe properties were taken as a volumetric average of the wire metal and polyimide insulation constituents and the copper plane regions were converted to a blended material that averaged the properties of FR-4 and copper by their approximate proportion in the thickness of the board. The solid-solid and solid-liquid transition for docosane were approximated as a single transition without hysteresis, having a latent heat of 252 J/g and beginning and ending melting temperatures of 41 °C and 44.5 °C. The underfilling of the bore was approximated by then setting the latent heat to 80% of its material value. Temperatures lower than the melting point were assumed to be sufficiently solid and above the highest melting point to be sufficiently liquid. Due to likely imperfections in the model, and non-idealities in the assembly, when assessing sufficiency, an additional safety factor was added (~5 °C).

Errors in the model are more likely to affect the ‘release’ than the independence behavior. The gripper release maxima are more dependent on the approximated and idealized geometries within a gripper, which are thin relative to the duration of a heat pulse. The independence behavior is more dependent on the overall gripper geometry, spacing, separating material and total energy of the heat pulse, which was shorter than the inter-gripper diffusion time. Further, release timing had been previously found for model devices, whereas this simulation mainly investigated a higher gripper count.

Heating periods of prescribed times were applied to sets of the heaters, and the maximum, minimum and average temperature were monitored in the PCM regions. Starting at a temperature of 20 °C, a model gripper could be fully melted after heating at a rate of 250 mW for 12.5 ms. At its warmest, the minimum temperature of the PCM reached 49.2 °C, occurring approximately 3 ms after the heating ended (Figure S4). From the beginning of the heating pulse to the time of the maximum temperature, the grippers had practically identical temperature rises and did not have a significant effect on each other; the number of grippers ‘on’ only changed the maximum by ~1 °C. This independence based on the short heat pulses and short distance between a micro-gripper’s heater and PCM relative to the distance between micro-grippers greatly simplifies the use of the microdrive. The number of grippers turned on did affect the cooling time for the gripper region over the one hundred millisecond time scale, with the effect being greater towards the center of the pattern.

**Figure S4.**
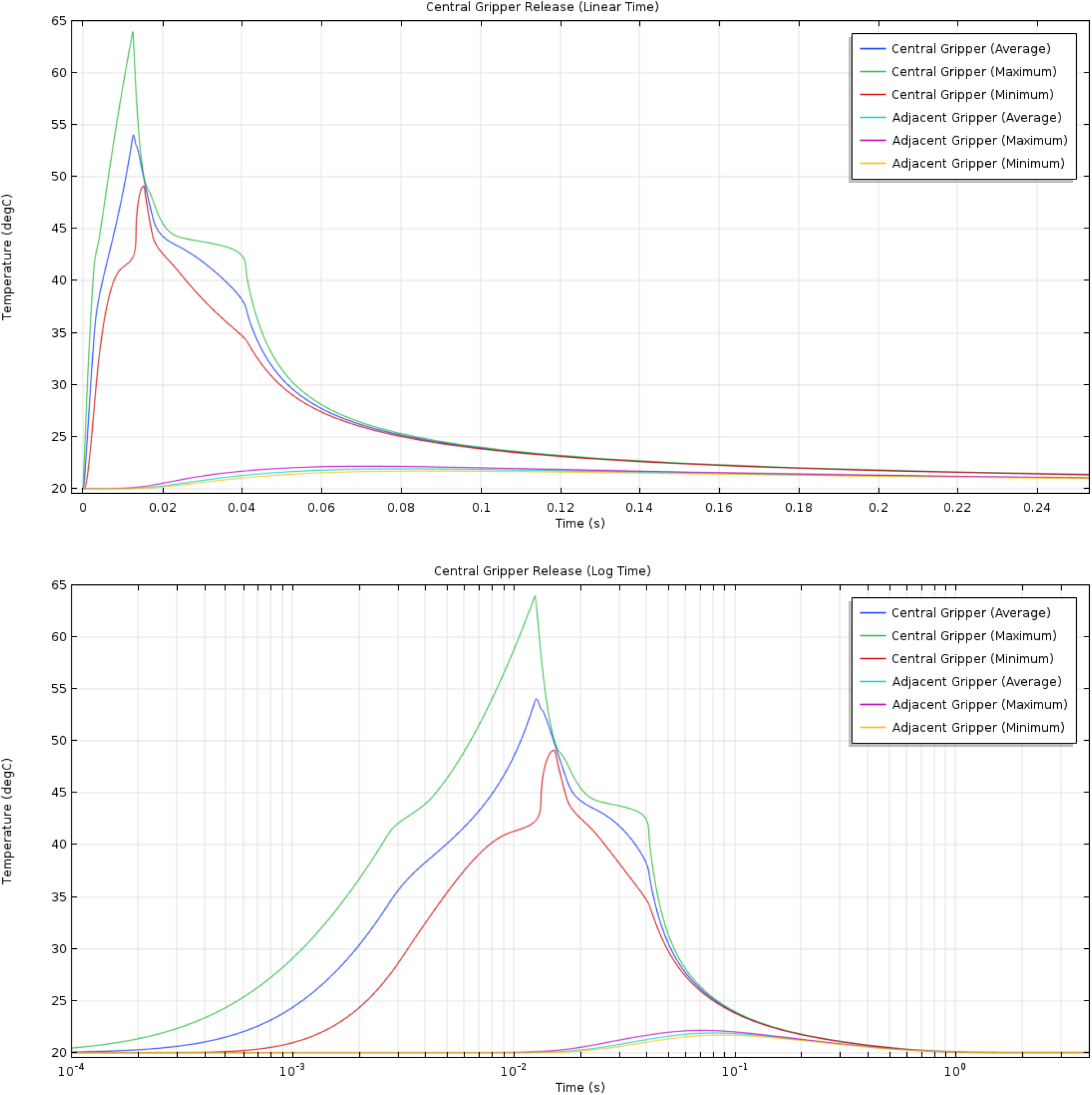
(Top) Linear and (Bottom) log time plots of the simulated average, maximum and minimum temperatures for the PCM in a single ‘on’ gripper and one adjacent to it

For independence, the worst-case thermal scenario is when the central gripper is ‘off’ but all of the surrounding 15 are ‘on’. Using the same 12.5 ms heating time, the center gripper reached a max temperature of 34.3 °C ~110 ms after the beginning of the heat pulse (Figure S5, Supplementary Video 7, 8). The temperature change in the central gripper indicates substantial inter-gripper thermal coupling, but the heating efficiency and phase change non-linearity enable independence.

**Figure S5.**
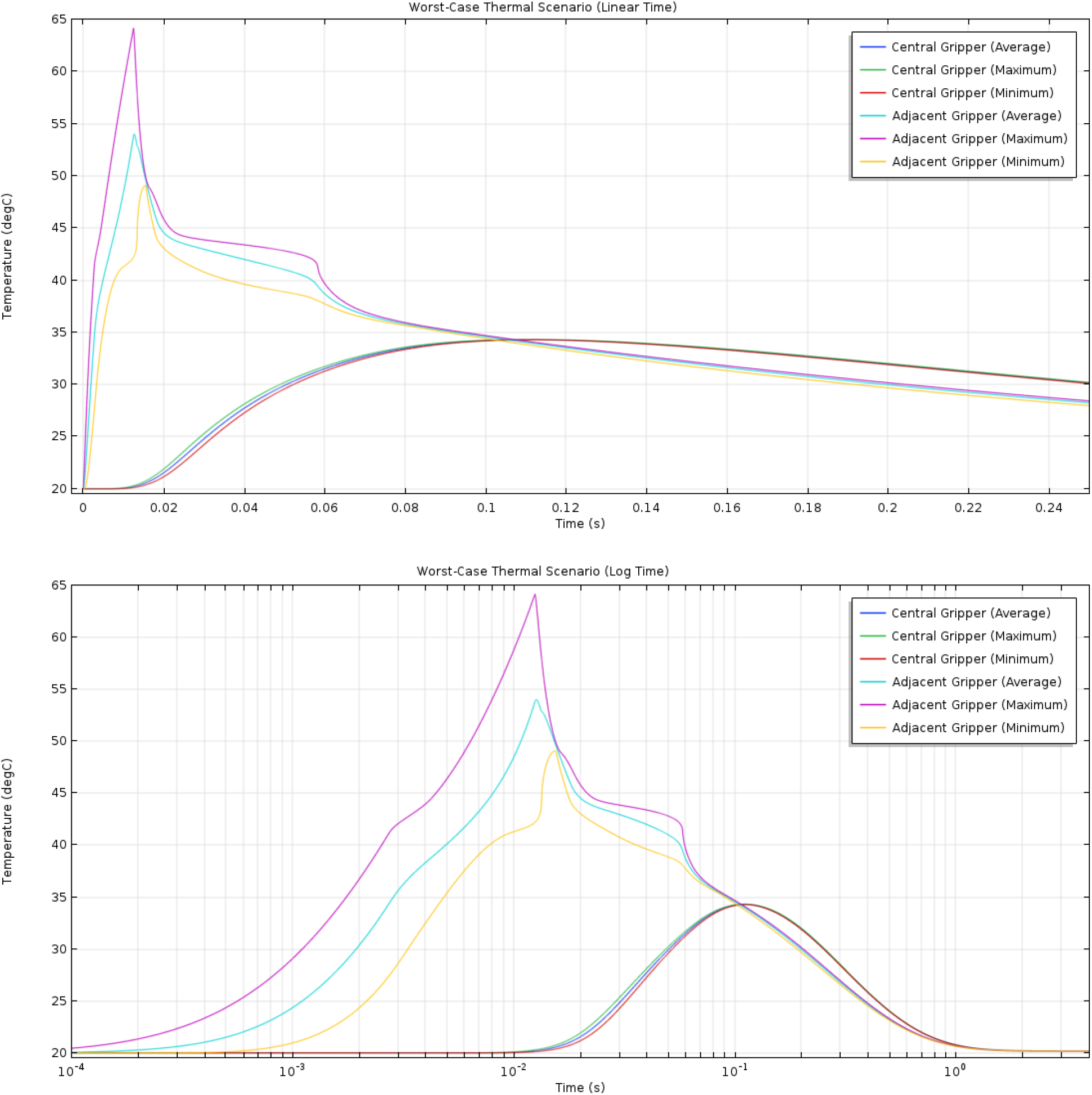
(Top) Linear and (Bottom) log time plots of the simulated average, maximum and minimum temperatures for the PCM of an ‘on’ gripper and the central ‘off’ gripper in the ‘worst-case’ scenario

### Detailed description of Tetrode Loading for Implantable Devices

For each probe, approximately 25 cm lengths of twisted wire tetrodes were prepared beforehand, with one end snipped and the other end with the wires still separated. Initially, the cage, microdrive and EICB were slightly separated and angled relative to each other such that the holes of the bottom cannula and top board were visible for threading. Under a microscope, the tetrodes were individually threaded through each via and stuck onto a piece of adhesive below the assembly to prevent it from withdrawing through the cannula. Throughout this process, tetrodes were only gripped by a section near the trimmed end that would later be trimmed away. Before threading another tetrode, two small particles of PCM were attached onto the tetrode near the boards, and then a heat lamp was used to melt them into beads. This process was repeated with all subsequent tetrodes. After all the tetrodes had been threaded, the cage, microdrive and EICB were aligned and secured.

Each tetrode wire was then individually connected to the EICB. This can be done in the standard crimp style, but we instead used a slightly modified method to reduce the risk of incompletely shearing the insulation or completely cutting through the wire. The insulation was burnt off the connection ends of the tetrodes using a small heating element, and then the ends were wrapped and tied around the crimp pins and adhered using conductive silver paint. The pins were then inserted into their crimp holes and stress was decoupled from the connection by applying wax over the pins.

The heating lamp was then used to heat the assembly and remelt the PCM on all the tetrodes. Each tetrode was pulled through the bottom cannula until only enough remained between the EICB vias and the crimp locations to allow for the desired range of motion. During this sliding process, the PCM beads were wicked into their bores such that they would grip the probe upon cooling. The ends of the tetrodes were snipped while the PCM was still melted in order to preventing any kinking during the cutting process.

The top cap of the EICB was then placed to protect the tetrode loops. Functionality of the tetrodes was checked by using the microdrive. If loaded successfully, there should be no kinks on the tetrodes, which would lead to bowing and poor movement behavior.

